# Plasticity in cryoprotectant synthesis involves coordinated shunting away from pyruvate production

**DOI:** 10.1101/2024.10.16.618698

**Authors:** Karin R. L. van der Burg, Yasmin Bozorgi, Katie Gyte, Amanda D. Roe, Katie E. Marshall

## Abstract

Insects living in temperate regions often accumulate a large amount of glycerol during winter to avoid freezing. This seasonal accrual of glycerol is generally produced from glycogen reserves through the pentose phosphate pathway. An alternative pathway to produce glycerol is through glycolysis, normally used for pyruvate production for eventual ATP synthesis. Aside from seasonal accumulation, some insects will also rapidly increase glycerol production as a short-term response to a sudden cold event, thereby increasing cold hardiness when necessary. In the eastern spruce budworm *Choristoneura fumiferana*, this plasticity in cold hardiness is locally adapted, where northern populations produce more glycerol upon cold shock. Here we investigate how glycerol is produced during the rapid plastic response to fluctuating cold conditions, and whether this pathway could be a target of local adaptation. After a period of repeated cold exposure, we found evidence of increased enzyme activity and increased mRNA abundance of several proteins associated with glycolysis, and a downregulation in expression of glucose-6-phosphate dehydrogenase, associated with pentose phosphate. Pyruvate production is prevented through downregulation of glyceraldehyde-3-phosphate dehydrogenase. We found higher overall enzyme activity and glycerol accumulation in a northern population from Alberta, although there was no evidence of an interaction effect between population and cold shock treatment. This is one the first studies to show a mechanistic basis of such plasticity in cold hardiness.

## Introduction

One of the biggest challenges faced by insects living in temperate regions is surviving harsh winters with limited food availability and extremely cold temperatures. In the North American boreal forest, temperatures can drop below −30 °C, and insects employ several strategies to cope with these extreme conditions (Storey and Storey, 2012). For example, many insects will enter a state of dormancy called diapause, characterised by low metabolic activity, arrested development and transcriptional shut-down (Denlinger, 2023). In addition, many insects will employ freeze avoidance mechanisms, such as synthesising antifreeze proteins, significant desiccation, and producing sugar or polyol cryoprotectants such as glycerol (Teets et al., 2023). Indeed, many species of insect accumulate extremely high concentrations of glycerol during winter months (Han and Bauce, 1995a; Rickards et al., 1987) that increase cold hardiness. Aside from seasonal accumulation of glycerol, insects can also employ short-term responses to sudden cold events. Upon cold exposure, cold hardiness can rapidly increase due to increased glycerol production (Butterson et al., 2021; Marshall and Sinclair, 2012). This plasticity in cold hardiness can be employed within minutes of sudden cold exposure, and allows insects to cope with thermal variability that is common during winters (Teets et al., 2020).

Because winter conditions can vary across regions, insects often show local adaptation in cold hardiness mechanisms (Marshall et al., 2020). Indeed, variation in diapause incidence over a latitudinal gradient has been described repeatedly in several species, and is one of the most robust examples of clinal variation in a phenotype (Demont and Blanckenhorn, 2008; Posledovich et al., 2015; Tyukmaeva et al., 2011). Cold hardiness also varies over a geographic range, where populations from more poleward regions tend to be more resistant to colder temperatures than populations from less cold regions (Sinclair et al., 2012). For example, survival after cold shock and recovery from a chill coma both increased with poleward latitude in *Drosophila melanogaster* (Hoffmann et al., 2002). Freeze avoidance mechanisms can also be locally adapted, for example in the rice stem borer (*Chilo suppressalis*) in Japan, northern populations accumulate more glycerol faster during winter months than compared to populations from the southwest (Ishiguro et al., 2007). Thus, species with large distributions are likely to have population-specific cold hardiness adaptations, although studies that investigate the physiological mechanisms underlying these local adaptations are relatively rare.

Cold hardiness measures are energetically expensive to produce, thus overwintering insects face significant energetic costs while at the same time food resources are limited and feeding is often avoided for months to avoid the accumulation of ice nucleators in the digestive tract (Hahn and Denlinger, 2011; Sinclair, 2015). The necessary cryoprotectants are synthesised from the limited supply of glycogen reserves accumulated prior to winter (Storey and Storey, 2012). Under normal, non-winter conditions, glycogen reserves would be used for the production of pyruvate and eventually ATP through the tricarboxylic acid cycle. However, during winter, insects are mostly dormant with suppressed metabolism, thus the production of excess ATP is wasteful and should be avoided. This means that insects must somehow shunt glucose equivalents away from pyruvate production and towards cryoprotectant polyol accumulation (Storey and Storey, 2012). In the case of glycerol production, this is broadly believed to be accomplished by reversible phosphorylation of glucose-6-phosphate dehydrogenase (G6PDH) and phosphofructokinase (PFK), which shunts glucose equivalents towards the pentose phosphate pathway and away from glycolysis. This has the advantage of producing NADPH, which is later needed for the production of glycerol (Storey and Storey, 2012). However, the use of this pathway has only been characterised for seasonal accumulation of glycerol rather than the short-term responses to cold that fluctuating cold conditions can induce. It is possible for glycerol to be synthesised instead from glycerol-3-phosphate, which is produced from dihydroxyacetone phosphate during glycolysis (Storey and Storey, 2012). This pathway is possibly faster, and thus more appropriate for the rapid plastic response necessary for a sudden exposure to cold temperatures. However, this pathway uses ATP, which has limited availability in winter, and downstream enzymes that catalyse reactions leading to pyruvate accumulation would need their activity to be reduced so that pyruvate production is not increased beyond the needs of an animal in deep metabolic suppression. In this study, we investigate which glycerol production pathway is used during the rapid plastic response to fluctuating cold conditions, and whether this pathway could be a target of local adaptation.

The eastern spruce budworm (*Choristoneura fumiferana*) is the most destructive insect defoliator of conifers in Canada’s boreal forest (Laurentian Forestry Centre, 2018). Eastern spruce budworm are adapted to survive harsh conditions by entering diapause during the second instar larval stage (Régnière, 1990; Han and Bauce, 1998). In this state, larvae suppress their supercooling point (SCP; the lowest temperature reached before freezing (Sinclair et al., 2015)) to temperatures as low as −40 °C through upregulation of cold hardiness mechanisms such as the increased production of glycerol and hyperactive antifreeze proteins (Han and Bauce, 1995b; Marshall and Roe, 2021; Butterson et al., 2021). As 2nd instar budworm larvae enter and progress into diapause, they accumulate circa 3 M glycerol (Han and Bauce, 1995b; Marshall and Sinclair, 2015). A genetic analysis across its range showed the species consists of three subpopulations: Western, Central and Eastern populations (Lumley et al., 2020). One of the differentiating SNPs between populations was located within glycerol-3-phosphate dehydrogenase, a gene associated with the glycolysis pathway (Storey and Storey, 2012). Indeed, high latitude Central populations accumulate more glycerol during diapause as compared to lower latitude Eastern populations (Butterson et al., 2021). Thus, spruce budworm populations have adaptive differences in cold tolerance where high latitude populations suppress their SCP more than lower latitude populations (Fig. 1A), meaning that high latitude populations are more cold tolerant than low latitude spruce budworm (Butterson et al., 2021). In addition to seasonal accumulation of glycerol, spruce budworm populations also show variation in the short-term response to cold exposure. Upon experiencing cold temperatures, Central populations collected from Inuvik and Alberta will rapidly accumulate more glycerol as compared to Eastern populations collected from northern Ontario and Quebec (Butterson et al., 2021). This plasticity in glycerol production allows for greater conservation in energy resources– glycerol accumulation is energetically costly, and production mechanisms are only employed when necessary, leaving more energy available for maintaining homeostasis. However, the precise mechanisms by which local adaptation in cold hardiness plasticity is regulated are currently unknown.

**Figure 1.**
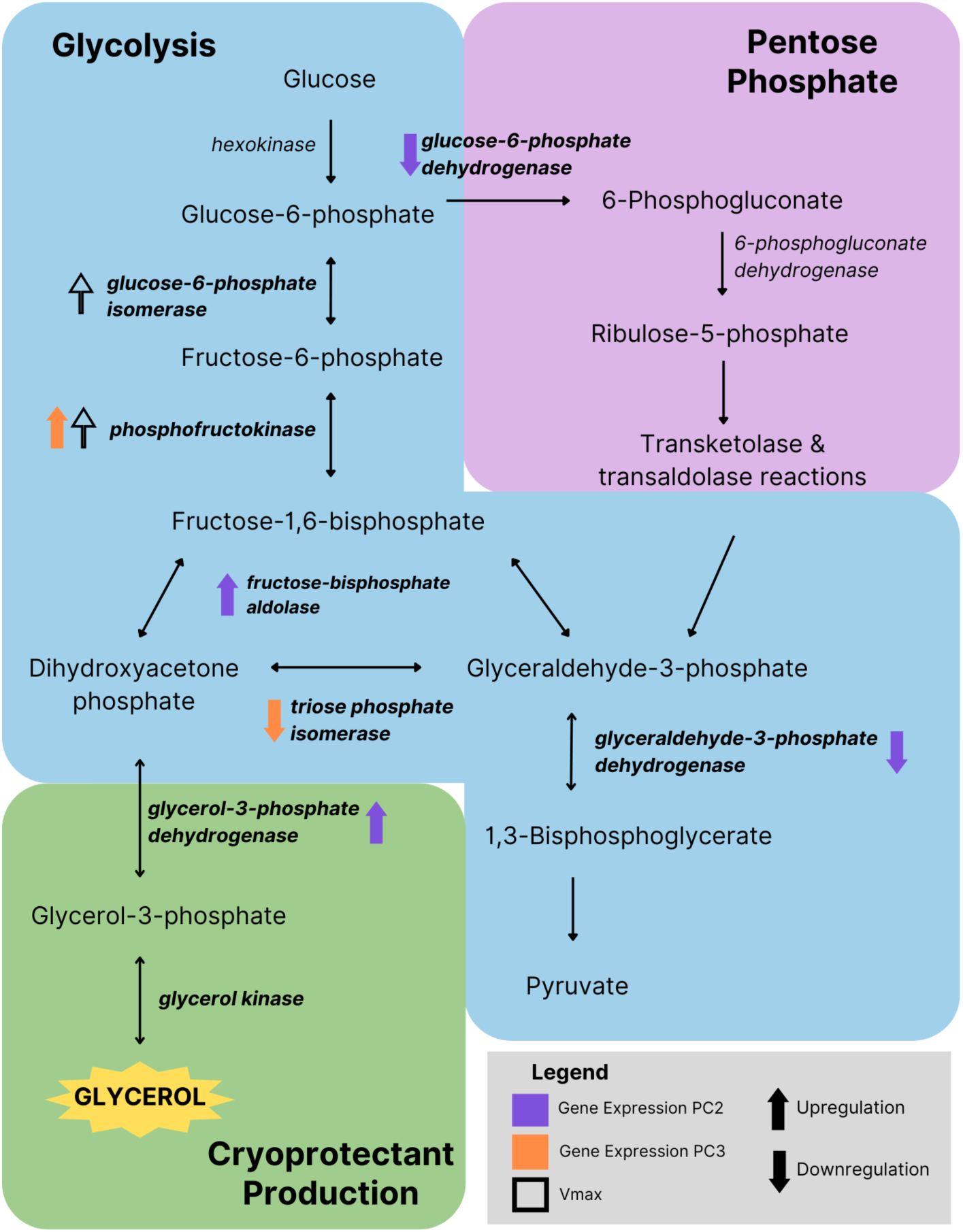
Metabolic pathways involved in glycerol production in *Choristoneura fumiferana*. Closed arrows represent change in principal component two and three of relative gene expression, and open arrows represent change in Vmax.

In this study, we investigate how glycerol is produced in response to these fluctuating conditions, and whether local adaptation in this rapid response pathway exists. We investigate the activity of key genes representative of either the pentose phosphate pathway or the glycolysis pathway in response to repeated cold exposure. We assay both expression and enzyme activity for glucose-6-phosphate dehydrogenase and glucose-6-phosphate isomerase, chosen because these enzymes are the first branchpoint between glycolysis and the pentose phosphate cycle. Additional genes represent different downstream steps in either pathway. By measuring both gene expression and enzyme activity we can investigate not only how the genetic processes change in response to environmental conditions, but also how that gets translated into changes in cellular metabolic processes. To test for local adaptation in glycerol production, we tested gene expression and enzyme activity in diapausing larvae that originated from Central and Eastern populations. We hypothesise that the pentose phosphate pathway is used for rapid response to cold exposure, as that pathway is more likely to prevent the production of unnecessary pyruvate. We would expect to see increased activity in gene expression and activity of enzymes associated with the pentose phosphate pathway upon repeated cold exposure of second instar spruce budworm larvae.

## Methods

### Sample preparation

All experiments were performed on second-instar diapausing eastern spruce budworm (*Choristoneura fumiferana*) from four different populations, all sourced by Insect Production and Quarantine Laboratory (IPQL) at the Great Lakes Forestry Centre in Sault Ste. Marie, Ontario, Canada (Perrault et al., 2021)Roe et al. 2018(Perrault et al., 2021). The four populations tested were the in-lab strain established in 1961 (Glfc:IPQL:Cfum, “IPQL” hereafter), and wild strains established in 2018 from Campellton, New Brunswick (Glfc:IPQL:CfumBNB01, “New Brunswick”), High Level, Alberta (Glfc:IPQL:CfumHAB, “Alberta”), and Inuvik, Northwest Territories (Glfc:IPQL:CfumINT01, “Inuvik”). Caterpillars were kept in an incubator held at 2°C during diapause and underwent experimental cold exposure conditions after 6-7 weeks in diapause. At this stage, caterpillars were aliquoted into microcentrifuge tubes with 45 larvae per tube, maintained at 2°C until time for use. Two experimental cold exposure conditions were tested: control and repeated cold exposure. The control group was held in a 2°C incubator for the duration of the cold exposure period. The repeated cold exposure (RCE) conditions were controlled using a milled aluminium block connected to a programmable refrigerated circulating bath. The RCE experimental group were subjected to five cycles of 2°C to −15°C temperature fluctuations, including 12 hours at 2°C followed by 12 hours of decreasing temperature to −15°C with a ramp rate of 0.05°C/min, holding for approximately 1 hour, and increasing back to 2°C to repeat the cycle. After 5 days of repeated cold exposure, samples were placed directly into a −80°C freezer until use.

### Metabolite and Enzymatic Assays

Samples were homogenised using 0.9 mm stainless steel beads in a Bullet Blender in a final volume of 900 µL homogenization buffer (20 mM Imidazole-HCl, 5 mM EDTA, 5 mM EGTA, 50 mM sodium fluoride, 0.1 mM PMSF, 0.5 mM DTT). Samples were centrifuged for 10 minutes at 15,000 × g at 4°C, then the supernatant was stored at −80 °C until use. Enzyme activity for three different glycolytic enzymes was measured in five separate biological replicates of 45 larvae each, and metabolite content was measured in each sample (15 biological replicates).

Free glycerol content was assayed by measuring absorbance of three technical replicates per sample at 540 nm in a SpectraMax M2 spectrophotometer (Molecular Devices, San Jose, USA) following the addition of Free Glycerol Reagent (MAK117, Sigma-Aldrich Canada Co., Oakville, Canada), and by using glycerol as a standard. We measured glucose-based glycogen content using a Type II glycogen from oyster standard (Millipore Sigma) and a phenol-sulphuric acid method following (Masuko et al., 2005), with absorbance measured at 490 nm. Finally, protein content was measured using a Bicinchronicinc acid kit (BCA1, Sigma-Aldrich Canada Co.) and bovine serum albumin as a standard (Millipore Sigma), with absorbance measured at 562 nm.

Enzyme activity assays were generally based on methods developed for another freeze avoiding lepidopteran (Rickards et al., 1987) and were conducted by measuring the consumption of NADH (phosphofructokinase) or the production of NADPH (glucose-6-phosphate isomerase and glucose-6-phosphate dehydrogenase) at 22 °C. Vmax and Km of each enzyme were estimated by running a series of substrate concentrations for each sample, following optimization of each assay. Glucose-6-phosphate dehydrogenase activity was measured across concentrations of glucose-6-phosphate (0, 0.02, 0.04, 0.08, 0.3, 1 mM) by adding assay mix (0.2 mM NADP+, 5 mM MgSO4, 20 mM imidazole-HCl buffer) with an NADPH standard curve (0, 0.005, 0.01, 0.025, 0.05, 0.1 mM) then monitoring the change in absorbance at 340 nm for 15 minutes (read every 30 s) (Fig S1). Glucose-6-phosphate isomerase activity was measured using a coupled reaction across concentrations of fructose-6-phosphate (0, 0.02, 0.04, 0.08, 0.3, 1 mM) by adding assay mix (0.2 mM NADP+, 5 mM MgSO4, 0.5 IU G6PDH, 20 mM imidazole-HCl buffer) with a NADPH standard curve (0, 0.005, 0.01, 0.025, 0.05, 0.1 mM) then monitoring the change in absorbance at 340 nm for 15 minutes (read every 30 s) (Fig. S2). Finally, phosphofructokinase activity was measured using a coupled reaction across concentrations of fructose-6-phosphate (0, 0.5, 2, 5, 20, 40 mM) by adding assay mix (4 mM ATP, 0.15 mM NADH, 50 mM KCl, 5 mM MgSO4, 0.5 IU aldolase, 0.5 IU triosephosphate isomerase, 2 IU G3PDH), 20 mM imidazole-HCl buffer) with a NADPH standard curve (0, 0.025, 0.05, 0.1, 0.15 mM) then monitoring the change in absorbance at 340 nm for 15 minutes (read every 30 s) (Fig. S3).

### Data analysis of enzyme assays

The effect of population origin and experimental condition on per larva metabolite concentration was modelled using general linear models with protein content as a covariate in the R language and environment (R Core Team, 2023). Absorbance data collected from the enzyme assays was converted into nmol NADPH produced (or consumed, in the case of PFK assays) using a standard curve. The reaction rate was defined as the rate of change in NADPH for each substrate concentration, then Michaelis-Menten curves were fitted using PAST4.04, which allowed the calculation of Vmax and Km of each enzyme assayed (Hammer et al., 2001). The effects of population origin and experimental condition on Vmax and Km were then modelled using general linear models in R.

### RNA Isolation and cDNA synthesis

We extracted RNA from 45-90 caterpillars using the ThermoFisher TRIzol™ Plus RNA Purification for RNA isolation according to the protocol. All samples were treated with a Qiagen On Column DNA Digestion to ensure there was no DNA contamination. Samples were eluted twice in 50µL of RNAse-free water. Concentration of RNA and contamination levels were quantified using a NanoDrop. cDNA synthesis was performed using qScript cDNA Synthesis Kit according to protocol. cDNA samples were diluted 10µL in 100µL water and stored at −20°C. For complete protocols, see SI.

### RT-qPCR assays

Gene selection was based on their relative position in the interrelated pathways of glycolysis and pentose phosphate (Fig 1). Genes at the junction between the two pathways were prioritised to investigate which pathway *C. fumiferana* is used for glycerol production in response to repeated cold exposure. Gene sequences from the *C. fumiferana* genome (Béliveau et al., 2022) were used to design primers (Table S1). For the reference genes, we used *Tubulin beta-1 chain* (*Tb1*) and *Ribosomal protein S15* (*RPS15)* based on their efficiency in previous studies (Lü et al., 2018; Zhou et al., 2018). Primer sequences were designed using primerQuest from IDT. All primers were designed to achieve *T_m_* values of 61°C and amplicon sizes of approximately 100bp. Real-time quantitative polymerase chain reaction (RT-qPCR) was conducted in 10µL volumes, containing 8.6µL of KiCqStart SYBR Green qPCR ReadyMix (Sigma), 0.4µL gene-specific primer, and 1µL cDNA template. Serial dilution standard curves were generated as follows: 1, 1:2, 1:4, 1:8. A Bio-Rad CFX96 C1000 Touch Real-Time PCR Detection system was used with the following protocol: 40 cycles at 95°C for 3 minutes, 95°C for 15 seconds, 56°C for 15 seconds, and 60°C for 1 minute. RT-qPCR reactions using the standard curve were performed in triplicates for all samples, except IPQL which was performed as a duplicate. No-template and no-reverse transcriptase controls were also used to identify any contamination. Cycle threshold (Ct) values were collected for all genes in each sample.

### Data analysis qPCR

Ct values of the triplicates (or duplicates) were averaged and used to calculate primer efficiencies. The slope of the regression between the log values of each serial dilution and average Ct values were calculated. Slope values were used to calculate primer efficiency based on the equation:

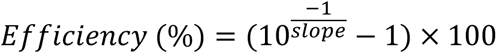

Since the calculated efficiencies were different across all genes (ranging from 83%-102%), the Pfaffl Method (Pfaffl, 2001) was used to analyse the data with increased reproducibility. Primer efficiency percentages were converted to amplification factors using the equation:

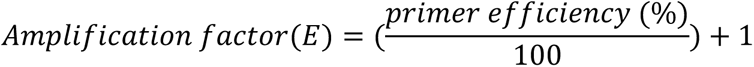

*Δ*Ct values were calculated by normalising raw Ct values to the IPQL control samples of each gene. Standard deviation calculations indicated RPS15 as the optimal housekeeping gene, and was therefore used in the Pfaffl formula. The Pfaffl formula was used to calculate the relative gene expression ratio:

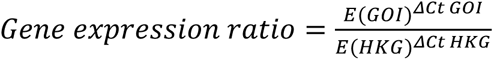

Average gene expression versus treatment (control or repeated cold exposure) in each population was calculated and plotted (see Sl Fig S4).

### Multivariate analysis

As our examined genes all were part of interconnected pathways that could be co-regulated, it was possible that there were correlations among the genes that could be modified by experimental conditions. Thus, we decided to test this possibility by conducting a principal components analysis (PCA) on the relative gene expression values to examine axes of variation (Fujisawa et al., 2021; Lenz et al., 2016). We then took the PC scores from this to generate new, orthogonal axes of variation, then used ANOVA to test whether population origin, experimental treatment, or their interaction significantly predicted PC score.

## Results

### Metabolites

We found that population origin and experimental condition had significant effects on metabolite content (Table 1; Figure 2). Generally, glycerol and glycogen content were significantly driven by both population origin and experimental condition. Glycerol content significantly increased following repeated cold exposure, and was significantly higher in the Alberta population (Figure 2A). Glycogen content also significantly increased following repeated cold exposure, and was generally higher in the wild origin populations (Alberta, Inuvik, and New Brunswick) than the IPQL population (Figure 2B).

**Table 1.** Table x. Effects of population, experimental condition (control or repeated cold exposure) and their interaction on metabolite content of diapausing *Choristoneura fumiferana* larvae. All reported values are F values from a two-way ANCOVA with protein content as a covariate, with those that are statistically significant (p < 0.05) bolded. N = 15 homogenates of 45 larvae per population × condition.

| Metabolite | Protein content | Population | Condition | Population x condition |
| --- | --- | --- | --- | --- |
| Free glycerol | **F<sub>1,111</sub> = 132.46** | **F<sub>3,111</sub> = 273.32** | **F<sub>1,111</sub> = 6.43** | F<sub>3,111</sub> = 0.18 |
| Glycogen | F<sub>1,111</sub> = 0.89 | **F<sub>3,111</sub> = 34.08** | **F<sub>1,111</sub> = 5.49** | F<sub>3,111</sub> = 0.59 |

**Figure 2.**
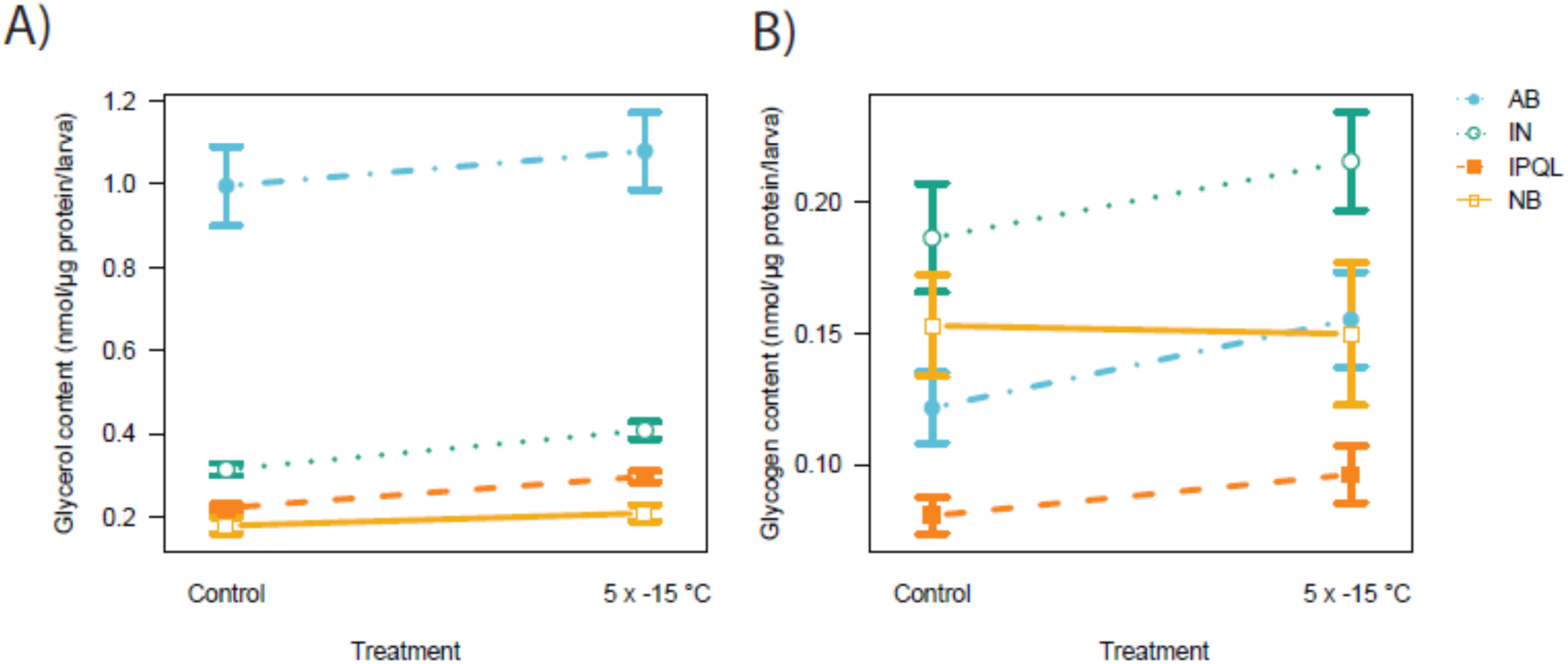
Metabolite content of diapausing eastern spruce budworm, *Choristoneura fumiferana*, varies as a function of both population origin and cold exposure (5 × −15 °C). A) Glycerol content (reported in nmol/µg protein/larva) varies significantly among populations as well as in response to repeated cold exposures. Populations tested include larval cultures sourced from northern Alberta (“AB”), Inuvik (“IN”), New Brunswick (“NB”) as well as from the standard laboratory colony (“IPQL”). B) Glycogen content (reported in nmol glucose/µg protein/larva) varies significantly among populations as well as in response to repeated cold exposures. All statistics reported in Table 1.

### Enzyme activity

We found that while there were significant effects of population origin on the Km of glucose-6-phosphate isomerase and glucose-6-phosphate dehydrogenase, there was no impact of repeated cold exposure on the Km of any of the enzymes assayed (Table 2, Figure 3). By contrast, repeated cold exposure significantly increased the Vmax of phosphofructokinase and glucose-6-phosphate isomerase (Table 3, Figure 4). In addition, Vmax was significantly affected by population origin in all three assayed enzymes (Table 3, Figure 4). However there was no consistent pattern of one particular population having higher enzyme activities or Km than any of the others.

**Table 2.** Effects of population, experimental condition (control or repeated cold exposure) and their interaction on the substrate binding affinity (Km) of three enzymes in diapausing *Choristoneura fumiferana* larvae. All reported values are F values from a two-way ANOVA, with those that are statistically significant (p < 0.05) bolded. N = 5 homogenates of 45 larvae per population × condition.

| Enzyme | Protein content | Population | Condition | Population x condition |
| --- | --- | --- | --- | --- |
| Glucose-6-phosphate dehydrogenase | $F_{1,31} = 0.11$ | **$F_{3,31} = 4.52$** | $F_{1,31} = 0.80$ | $F_{3,31} = 0.33$ |
| Glucose-6-phosphate isomerase | $F_{1,31} = 0.57$ | $F_{3,31} = 0.67$ | $F_{1,31} = 2.04$ | $F_{3,31} = 1.03$ |
| Phosphofructokinase | **$F_{1,31} = 35.80$** | **$F_{3,31} = 3.18$** | $F_{1,31} < 0.01$ | $F_{3,31} = 0.29$ |

**Figure 3.**
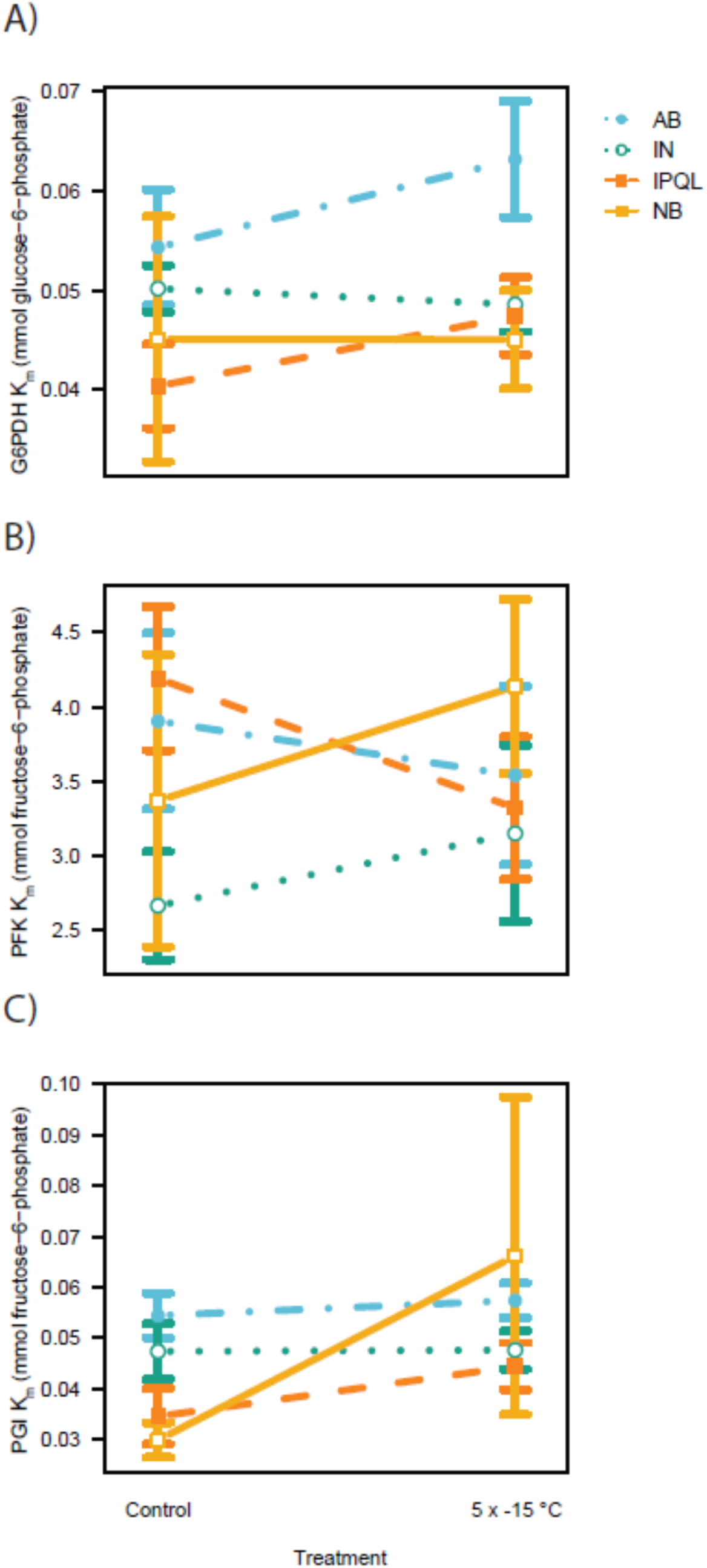
Substrate binding affinity (Km) of enzyme activities of three glycolytic enzymes in diapausing *Choristaneura fumiferana* larvae varies as a function of population origin, but not in response to repeated cold exposures (5 × −15 °C. A) Glucose-6-phosphate dehydrogenase binding affinity for glucose-6-phosphate varies significantly as a result of population origin. B) Phosphofructokinase binding affinity for fructose-6-phosphate varies as a function of population origin but not experimental treatment. C) Glucose-6-phosphate isomerase binding affinity for glucose-6-phosphate does not vary in response to condition or population. All statistics reported in Table 2.

**Table 3.** Effects of population, experimental condition (control or repeated cold exposure) and their interaction on the maximal turnover rate (Vmax) of three enzymes in diapausing *Choristoneura fumiferana*. All reported values are F values from a two-way ANOVA, with those that are statistically significant (p < 0.05) bolded. N = 5 per population × condition.

| Enzyme | Protein content | Population | Condition | Population x condition |
| --- | --- | --- | --- | --- |
| Glucose-6-phosphate dehydrogenase | **$F_{1,31} = 8.61$** | **$F_{3,31} = 8.06$** | $F_{1,31} = 0.63$ | $F_{3,31} = 0.25$ |
| Glucose-6-phosphate isomerase | **$F_{1,31} = 108.00$** | **$F_{3,31} = 15.92$** | **$F_{1,31} = 17.32$** | $F_{3,31} = 1.91$ |
| Phosphofructokinase | **$F_{1,31} = 61.58$** | **$F_{3,31} = 28.45$** | **$F_{1,31} = 15.58$** | $F_{3,31} = 0.55$ |

**Figure 4.**
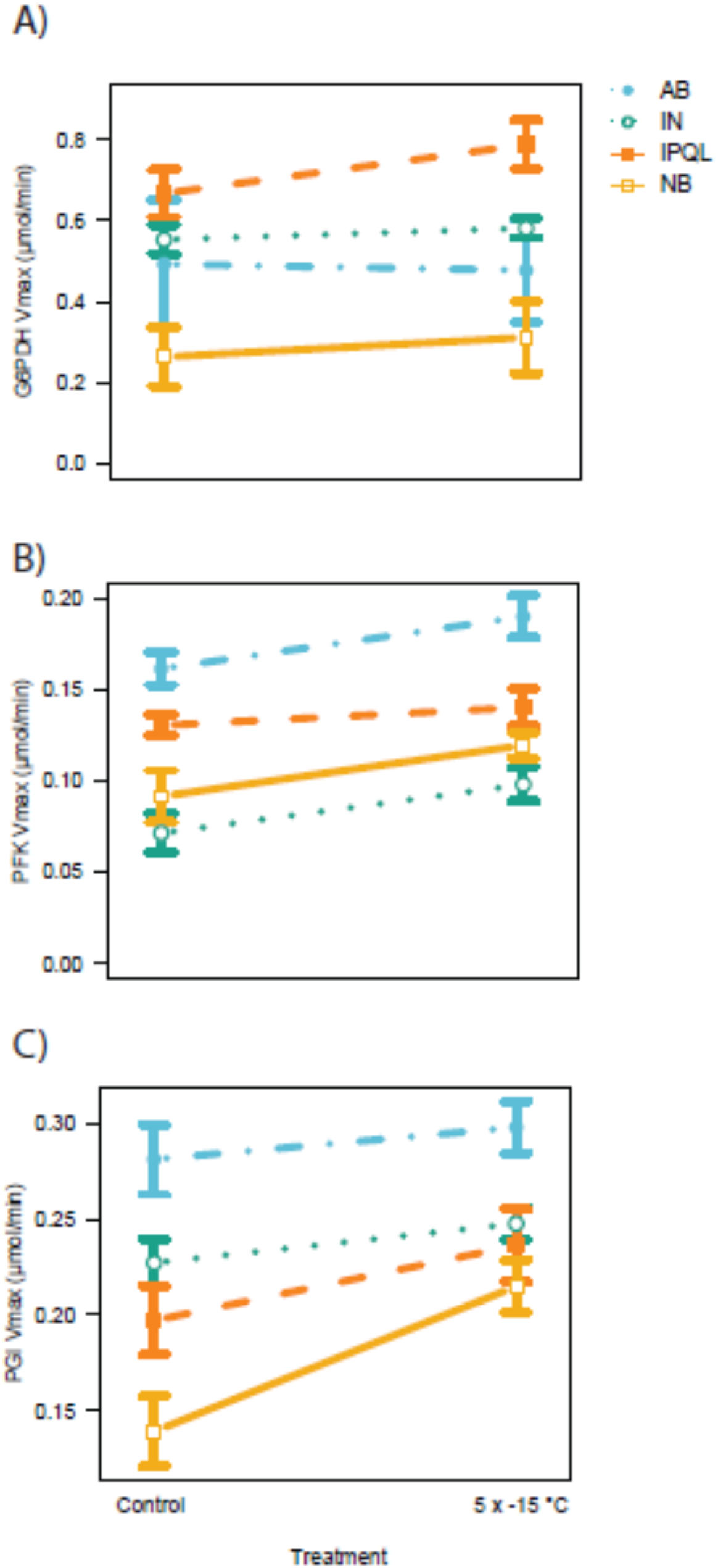
Maximal turnover rate (Vmax) of enzyme activities of three glycolytic enzymes in diapausing *Choristaneura fumiferana* larvae varies as a function of population origin and experimental treatment. A) Glucose-6-phosphate dehydrogenase maximal activity only varies in response to population origin B) Phosphofructokinase Vmax varies as a function of population origin and experimental treatment. C) Glucose-6-phosphate isomerase Vmax varies in response to both experimental condition and population. All statistics reported in Table 3.

### Multivariate analysis qPCR data

Our PCA indicated that three principle components could explain most of the variation in the gene expression data (a total of 82.94% of the variation, Table 4). PC1 (45.79% of the variation) appeared to describe an axis of greater or lesser overall gene expression, as the expression of all genes loaded significantly on this axis. PC2 (22.4% of the variation) had four genes that loaded significantly: glyceraldehyde-3-phosphate dehydrogenase and glucose-6-phosphate dehydrogenase both loaded positively while fructose-3-phosphate-aldolase and glycerol-3-phosphate dehydrogenase both loaded negatively. Finally, PC3 (14.8% of the variation) contained three highly-loaded genes: glucose-6-phosphate isomerase and phosphofructokinase loaded positively, while triose phosphate isomerase loaded negatively.

**Table 4.** Loadings of the expression of each gene on each major principal component. Loadings with an absolute value of greater than 0.25 are bolded.

| Gene | PC1 | PC2 | PC3 |
| --- | --- | --- | --- |
| Triose phosphate isomerase | **-0.341** | -0.117 | **0.527** |
| Glyceraldehyde-3-phosphate dehydrogenase | **-0.388** | **0.340** | 0.114 |
| Glucose-6-phosphate dehydrogenase | **-0.259** | **0.692** | -0.045 |
| Glucose-6-phosphate isomerase | **-0.404** | 0.013 | -0.071 |
| Fructose-3-phosphate aldolase | **-0.414** | **-0.283** | -0.099 |
| Glycerol-3-phosphate dehydrogenase | **-0.340** | **-0.554** | -0.061 |
| Glycerol kinase | **-0.422** | 0.061 | 0.093 |
| Phosphofructokinase | -0.195 | 0.015 | **-0.825** |
| Percent variance explained | 58.44% | 15.25% | 13.84% |

We then examined how population origin, experimental treatment, and their interaction influenced PC1, PC2, and PC3 scores using ANOVA. Population origin significantly predicted PC1 score (F_3,15_ = 12.38, p < 0.001), but neither cold exposure nor the interaction between population and cold exposure had a significant effect (p > 0.9 in both cases). While there was no significant interaction between population origin and cold exposure on PC2 score (F_3,15_ = 1.36, p = 0.292), there was a significant effect of population origin (F_3,15_ = 3.32, p = 0.049) whereby the New Brunswick population had significantly higher PC2 values. Given the low power of the experimental design to test interactive effects, but that plotting of the data suggested that the IPQL population did not respond to repeated cold exposure the way the other populations did, we tested the effect of dropping the IPQL population from the analysis. This caused the effect of repeated cold exposure to be statistically significant (F_3,15_ = 5.38, p = 0.039), whereby repeated cold exposure led to lower PC2 scores (i.e. higher glycerol-3-phosphate dehydrogenase and fructose-3-phosphate aldolase expression and lower glucose-6-phosphate dehydrogenase and glyceraldehyde-3-phosphate dehydrogenase expression; Figure 5, Table 4). Finally, both population origin (F_3,15_ = 6.35, p = 0.024) and repeated cold exposure (F_3,15_ = 14.10, p < 0.001) significantly drove PC3 score whereby repeated cold exposure led to lower PC3 scores (i.e. higher expression of phosphofructokinase and lower expression of triose phosphate isomerase).

**Figure 5.**
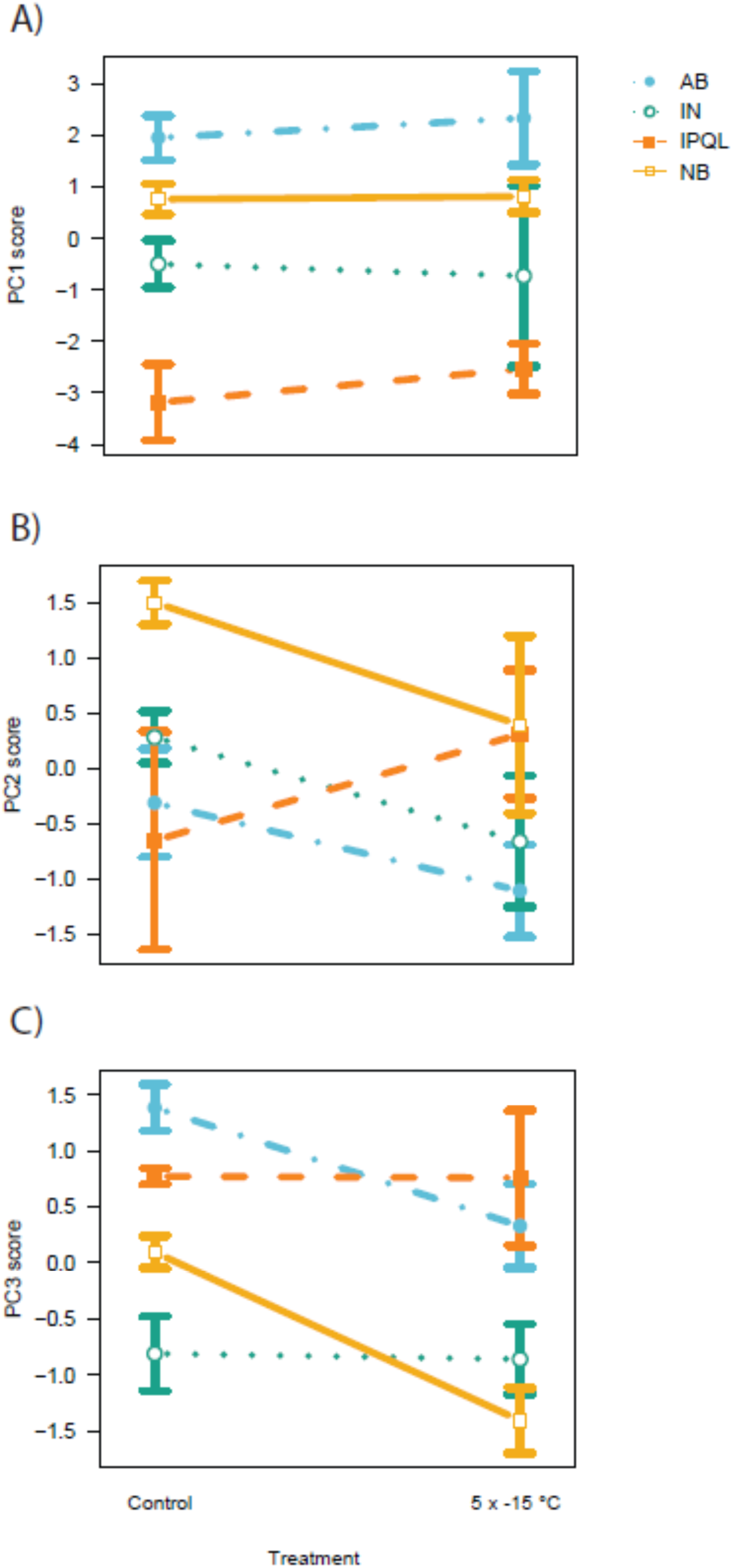
PCA analysis indicated that three principal components explain most of the variation in the expression data. A. PC1 showed a population origin effect, and appeared to describe overall gene expression. B. PC2 showed a population origin affect, where New Brunswick had significantly higher loading. C. Repeated cold exposure leads to lower PC3 scores.

## Discussion

Cold hardiness is a crucial mechanism through which insects can survive sub-zero temperatures, yet the precise mechanisms through which this hardiness is achieved is still unknown. Here, we investigated the mechanistic basis of glycerol production in response to repeated cold exposure in the eastern spruce budworm. We hypothesised that upon cold exposure, spruce budworm would shunt glycogen through the pentose phosphate pathway, as this is the pathway used for slow accumulation of glycerol throughout the season (Han and Bauce, 1995a), and would prevent the production of unnecessary pyruvate (Storey and Storey, 2012). Contrary to expectation, we found an upregulation of several transcripts and increased enzyme activity of proteins associated with glycolysis (Fig 4), indicating that glycolysis is the preferred pathway for mounting a rapid response to cold exposure in the eastern spruce budworm.

We found that glucose-6-phosphate dehydrogenase, an enzyme early in the pentose phosphate pathway, was downregulated following repeated cold exposure, indicating that this pathway is likely not used for rapid accumulation of glycerol. Pyruvate production is likely prevented through the downregulation of expression of both triose phosphate isomerase and glyceraldehyde-3-phosphate dehydrogenase, which reduces flux towards pyruvate. When put together, this data provides support for the use of the glycolysis pathway to rapidly respond to repeated cold exposure. Our results supporting glycolysis for rapid response glycerol production in response to repeated cold exposure is in direct contrast with several studies showing an increase of pentose phosphate flux upon oxidative stress in yeast, (Ralser et al., 2007), hypothermia in rats (Gallagher et al., 2009), and cold-hardening in gall moths (Muise and Storey, 1997). Gall moths showed reduced enzymatic activity for fructose-1,6-bisphosphatase during late fall, which would decrease glycolysis and possible recycling of glyceraldehyde 3-phosphate back to glycogen (Muise and Storey, 1997). The use of pentose phosphate under these stressful conditions is thought to be beneficial due to the increase in production of NADPH, an antioxidant that can protect cells against an increase in radical oxygen species (Ralser et al., 2007), and/or the demand for NADPH during downstream glycerol production. Although we do not have direct evidence of enzymatic activity of fructose-1,6-bisphosphatase, we show increased enzymatic activity and mRNA levels of two other enzymes associated with glycolysis upon repeated cold exposure. We would like to point out that pentose phosphate flux was still the dominant carbon flux pathway, as overall glucose-6-phosphate dehydrogenase activity (as measured by Vmax) was the highest of all enzymes investigated. This would indicate high overall activity of the pentose phosphate pathway, but this is not the pathway involved in mounting a rapid response to a repeated cold exposure. The seasonal accumulation of glycerol is a gradual process, where the use of the pentose phosphate pathway provides increased control, and an increase in redox power needed for glycerol production (and antioxidants). On the contrary, when a rapid response is necessary, glycolysis might be preferred despite the use of precious ATP and the risk of overproduction of unnecessary pyruvate.

We hypothesised that Central populations from higher latitudes would show evidence of higher enzymatic activity relating to glycerol synthesis upon cold exposure, as this would be consistent with our previous results; Central populations have higher induced glycerol accumulation relative to lower latitude populations (Butterson et al., 2021). We do see an overall population effect of both enzyme activity and mRNA abundance; where Alberta (a Central population) in particular had high enzyme activity and unusually high glycerol content (but not mRNA levels), and New Brunswick (an Eastern population) showed the lowest mRNA abundance. The interaction between population and repeated cold exposure was not significant, likely because our dataset was underpowered. Overall, we found evidence for some local adaption in glycolysis pathway activity.

A previous investigation of glycerol abundance in diapausing spruce budworm showed a strong population-specific effect, where Central populations had more glycerol overall, and a strong plastic response in glycerol accumulation upon cold exposure (Butterson et al., 2021). Surprisingly, we found a large amount of glycerol only in the Alberta population. We conducted our assays with all populations represented on the same 96-well plate, and on the same homogenate, and therefore believe that this high glycerol content is not the result of technical error, but represents a real difference among our populations. We also found that overall enzyme activity was higher in budworm from Alberta compared with Inuvik, although there was no significant effect of treatment. These results were surprising, as we would have expected similar levels of enzyme activity and glycerol content for both Inuvik and Alberta, and we would have expected a treatment effect. Our Inuvik results could be due to a low sample size, or an extended time of this population under laboratory conditions as compared to the previous experiment in (Butterson et al., 2021). The hypothesis that extended time in the laboratory has an effect on glycerol accumulation and plasticity has merit because the budworm from the IPQL population had the most divergent results, and the least plastic response to cold exposure. This stock was established in 1961, and had its last wild-type infusion around 40 years ago, and has thus been under laboratory conditions for much longer compared to other budworm stock populations (Perrault et al., 2021). Alternatively, our results for IPQL could be because it was established from an Eastern population from Northern Ontario, and thus has no need for a strong plastic response, but it is equally likely that drift and adaptation to laboratory conditions have had a significant effect on the cold-hardiness properties of this stock.

Genetic variation and local adaptation in glucose metabolism pathways in response to a warmer climate have been shown on several occasions, specifically for the gene glucose-6-phosphate isomerase (sometimes called phosphoglucose isomerase or PGI). Allelic variation in this gene varies along a climatic gradient, and has been shown to influence several life history traits in insects (Hill et al., 2021). Genetic variation in this gene has been directly linked to a change in enzymatic efficiency, especially under warmer temperatures (ie. 40°C, (Dahlhoff and Rank, 2000)). Interestingly, although we did test for variation in mRNA of glucose-6-phosphate isomerase, we did not see a significant change in expression of this gene, nor did we see a change in efficiency (Km). We did see a population and a treatment effect for capacity (Vmax) of this enzyme, but we also did for the other enzymes tested here. Thus, it appears that glucose-6-phosphate isomerase is not a target for adaptation in the eastern spruce budworm. It is of course possible that there is genetic variation across populations for this gene that we missed because we did not sequence our genes. It is also possible that there is a temperature-specific effect on adaptation, where below a certain temperature adaptation starts to target different genes. Previous results on glucose-6-phosphate isomerase show enzyme activity variation at temperatures ranging from 5 to 40°C, whereas we are studying enzyme activity variation after exposure to temperatures as low as −15°C. These results highlight the importance of investigating climate adaptations to different temperatures and in different species, as adaptation can take many forms.

This study investigates both mRNA levels as well enzymatic capacity (Vmax) and efficiency (Km) of different enzymes associated with glucose metabolism. The treatment of our qPCR data with a principal component analysis is unusual, but is appropriate because it allows us to account for individual variation between our samples. Also, the two orthogonal data types presented – enzymatic activity and quantitative mRNA PCR – both indicate increased glycolysis activity, providing further support for the validity of our results. Interrogating both mRNA and enzymatic processes allows us to investigate not only gene expression levels, but also the effect of post-translational modifications on enzyme activity. Previous results investigating glycerol production for cold hardiness showed fine-tuning of enzyme activity in glycolysis through reversible protein phosphorylation, a post-translational modification that changes enzyme efficiency (Storey and Storey, 2012). Interestingly, we did not see a change in enzyme activity (Km), indicating that fine-tuning through phosphorylation of enzymes is not likely to be part of the rapid response pathway to cold exposure. We did see an increase in Vmax, which indicates a higher capacity of the reaction, possibly due to a higher protein concentration. This higher concentration is likely a direct result of the higher gene expression we observed. Although a higher capacity in these enzymes almost certainly means that more glycerol is produced, we cannot know for certain how much glycogen is shunted through glycolysis. This is because Vmax describes the maximum capacity of the reaction rate *in vitro*, but not the actual rate *in vivo*. For that, we would need to measure the movement of radio-labelled glucose, which is an impossibility given that budworm do not feed between hatch and overwintering, and that injection into 100 ug animals with thick cuticles and 30% water content without significant damage is technically very challenging.

In conclusion, we show evidence of a mechanism for local adaptation and plasticity in cold tolerance in different spruce budworm populations across the boreal forest. Overall, we found that enzyme activity differs among populations, and can rapidly change in response to repeated cold exposure. While many studies show some form of phenotypic plasticity in response to extreme conditions, this is one the first studies to actually show a mechanistic basis of such plasticity. Our data aligns with previous studies showing local adaptation in glycerol production plasticity and evolution of glycerol associated genes, although we are the first to show a rapid response to repeated cold exposure through glycolysis. Future work will investigate in more depth the genetic basis of local adaptation in cold tolerance in this forest species.

## Data availability statement

All code and data are available on the Open Science Framework: https://osf.io/tbdge/?view_only=fd4b77134e1f4d8ba03530720a3013f3.

## Acknowledgments

We wish to express gratitude to everyone who contributed to this manuscript. Thanks to Ashlyn Wardlaw, Kerry Perrault, Alice (Yeuhong) Liu and the Insect Production team at the Great Lakes Forestry Centre for maintaining the spruce budworm colonies that were the focus of this project. KEM is supported by an NSERC Discovery Grant. KB is supported by the NSF award DBI-2208932; ADR was supported by Natural Resources Canada and the Healthy Forest Partnership through the Spruce Budworm Early Intervention Strategy - Phase II program financed by the Government of Canada in partnership with participating Provinces and forest industry.

