## Supplemental Information for "Plasticity in cryoprotectant synthesis involves coordinated shunting away from pyruvate production"

### Enzyme activity

#### Data analysis

Absorbance data collected from the enzyme assays was converted into nmol NADPH produced (or consumed, in the case of PFK assays) which was then plotted in a scatter graph in Microsoft Excel, showing change in NADPH over time. The rate of change in NADPH for each substrate concentration was determined using the linear trendline function in Excel, which corresponds to the reaction rate (Figs. S1-3). This substrate concentration vs. rate data was fitted as a Michaelis Menten (MM) curve using the data graphing software, PAST4.04. The MM curves generated using PAST also gave the Vmax and Km of each enzyme assayed (Figs. 2-3), which was subjected to ANOVA tests to determine the statistical significance of results.

### RT-qPCR sample prep

#### RNA Isolation

All steps were performed at room temperature (20-25 degrees celsius). RNAZap was used to remove contamination from all work surfaces (i.e., bench, fume hood) and any other items used during purification. The ThermoFisher TRIzol™ Plus RNA Purification was used to carry out RNA isolation on different tissue samples. 500µL of Trizol Reagent was added and incubated for 5 minutes to ensure dissociation of nucleoproteins. Samples were then disrupted for 5 minutes using homogenization beads and a bullet blender at high speed. Solution was separated from beads and 100µL of chloroform was added for lysis. Samples were then placed in the centrifuge for 15 minutes at 12,000 x g at 4 degrees celsius. Approximately 300µL of colorless upper aqueous phase (containing RNA) was transferred to a new tube, 300µL of 70% ethanol was added, and mixture was vortexed.

Samples were then transferred to spin cartridges and centrifuged at 12,000 x g for 15 seconds. Flow-through was discarded, spin cartridges were reinserted into collection tubes, and the process was repeated. Samples were treated with a Qiagen On Column DNA Digestion to ensure there was absolutely no DNA contamination. DNase I stock solution was made by combining 550µL of RNase-free water into DNase I vial and then was divided into 20µL aliquots (two samples per tube) and stored at -20 degrees. Approximately 140µL of Buffer RDD was added (70µL per sample), inverted to mix and briefly centrifuged.

To wash RNA on the membrane, 625µL of Wash Buffer I was added to the spin cartridge and centrifuged at 12,000 x g for 15 seconds. The flow-through was discarded and the spin cartridge was reinserted to the same collection tube. Next, 500µL of Wash Buffer II is added to the spin cartridge and centrifuged at 12,000 x g for 15 seconds. Flow-through was discarded and the process of adding Wash Buffer II was repeated. Samples were centrifuged at 12,000 x g for 1 minute to dry the membrane. Collection tube was discarded and the spin cartridge was inserted into a recovery tube. RNase-free water (1 x 50µL) was added to the center of the spin cartridge and incubated for 1 minute. Samples were then centrifuged at 12,000 x g for 2 minutes. Flow through was discarded, samples were reapplied to the column, and the process was repeated once more. Concentration of RNA and contamination levels were quantified using NanoDrop.

**cDNA synthesis**

cDNA synthesis was performed using qScript cDNA Synthesis Kit and all components were thawed, mixed, and centrifuged before use. The following were added to a PCR tube for each sample: 100ng RNA, nuclease-free water (variable quantities), 4µL qScript Reaction Mix (5X), and 1µL qScript RT. Solution was gently vortexed and briefly centrifuged. Tubes were then placed into a thermal cycler programmed: 22 degrees celsius for 5 minutes, 42 degrees for 30 minutes, 85 degrees for 5 minutes, 4 degree hold. cDNA samples were diluted 10μL in 100μL water and stored at -20 degree celsius.

**Table S1. Primer design**

| Primer Name | Primer Sequence (Forward) | Primer Sequence (Reverse) | Description |
|---|---|---|---|
| SBW-Tb1-2 | GTCTGGGTACTCTTCTCTGATT | GCCTACAAGGTTTCCAACTTAC | Tubulin beta-1 chain; sequence, housekeeping gene |
| SBW-RPS15-2 | GGCTTGTATGTAACGGAGAAC | GCTCCATTGTCGGCATTTA | Ribosomal protein S15; sequence, housekeeping gene |
| SBW-gly | GGACAGGAAGGGATTTGTTTC | CACCCTATTATCTCGGCCTATAA | Glycerol 3-phosphate dehydrogenase |
| SBW-gla3p | GCTATCAAGCAGAAGGTGAAG | GAGTGAGTGTCTCCGATGAA | Glyceraldehyde-3-phosphate dehydrogenase |
| SBW-g6pdh | TTTCGGAAGTCCGAGAGAAA | GAACCCGCAAGATATTCGTTAG | Glucose-6-phosphate dehydrogenase |
| SBW-g6p | AGCGTCTGAACCCTGAAA | GAACCAGTTCTTAGCTGATGTG | Glucose-6-phosphate isomerase |

| | | | |
|---|---|---|---|
| SBW-fba | TCGAGAACACTGAGGAGAAC | AGTCTCGTGGAAGAGGATTAC | Fructose bisphosphate aldolase |
| SBW-6p6k | CTCGAACTTCAGCCACAAAC | CCATAGCAGGCGAGAGTAAT | 6-phosphofructo-6-kinase |
| SBW-triop | GGTGGTAACTGGAAGATGAATG | CTCTTGACATAGGCCAGATAGA | Triose phosphate isomerase |
| SBW-gkinase | TAGCAGTTGGCGTTACAAATC | CCTCATATCCAGCCACACTAT | Glycerol kinase |

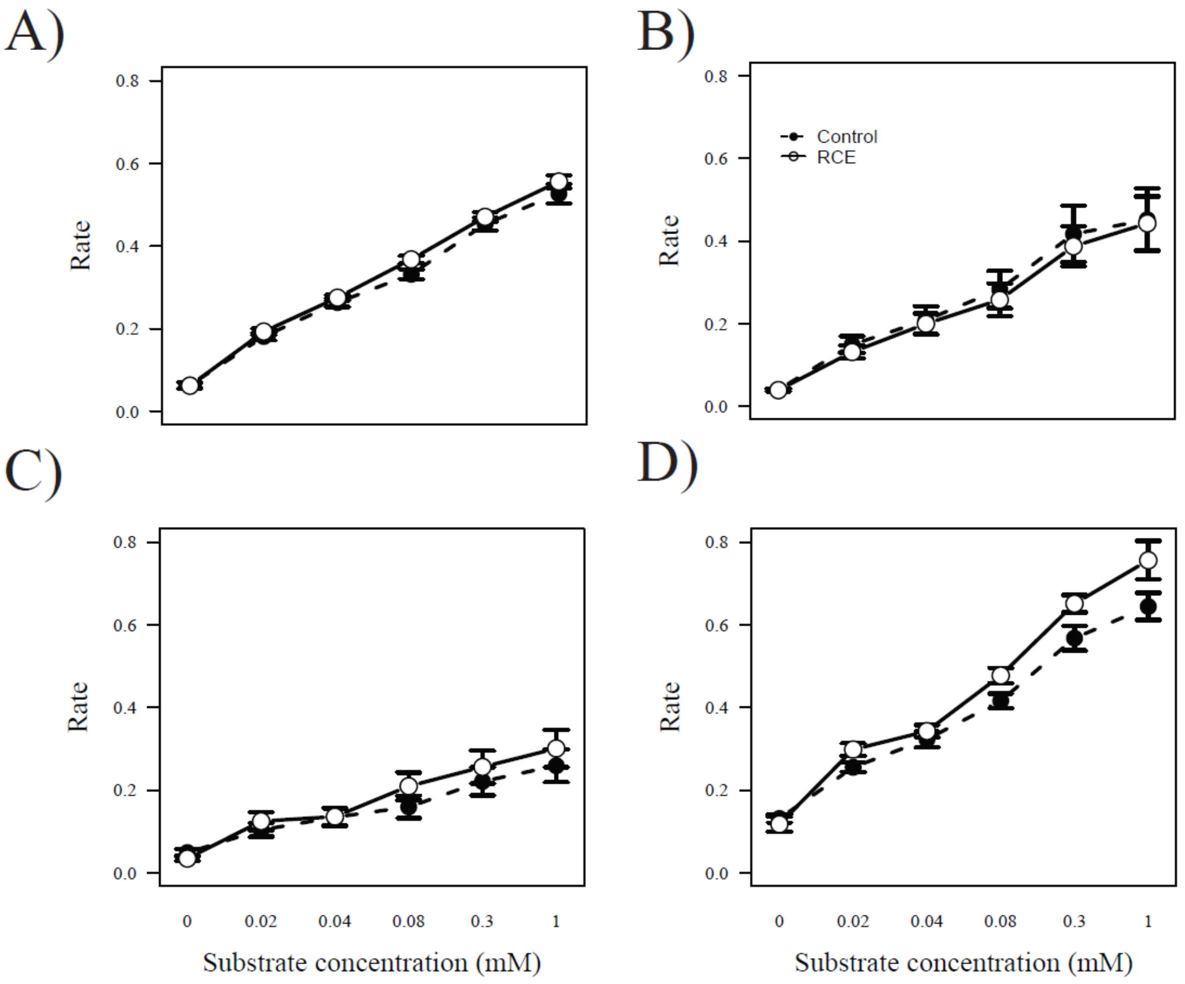


SI Figure 1: Reaction rate (nmol NADPH produced per minute) of glucose-6-phosphate dehydrogenase in *Choristoneura fumiferana* across populations generally does not increase following repeated cold exposure (RCE) across substrate concentrations. A) Inuvik population, B) Alberta population, C) New Brunswick population, and D) IPQL strain. N = 5 biological replicates per cold exposure type.

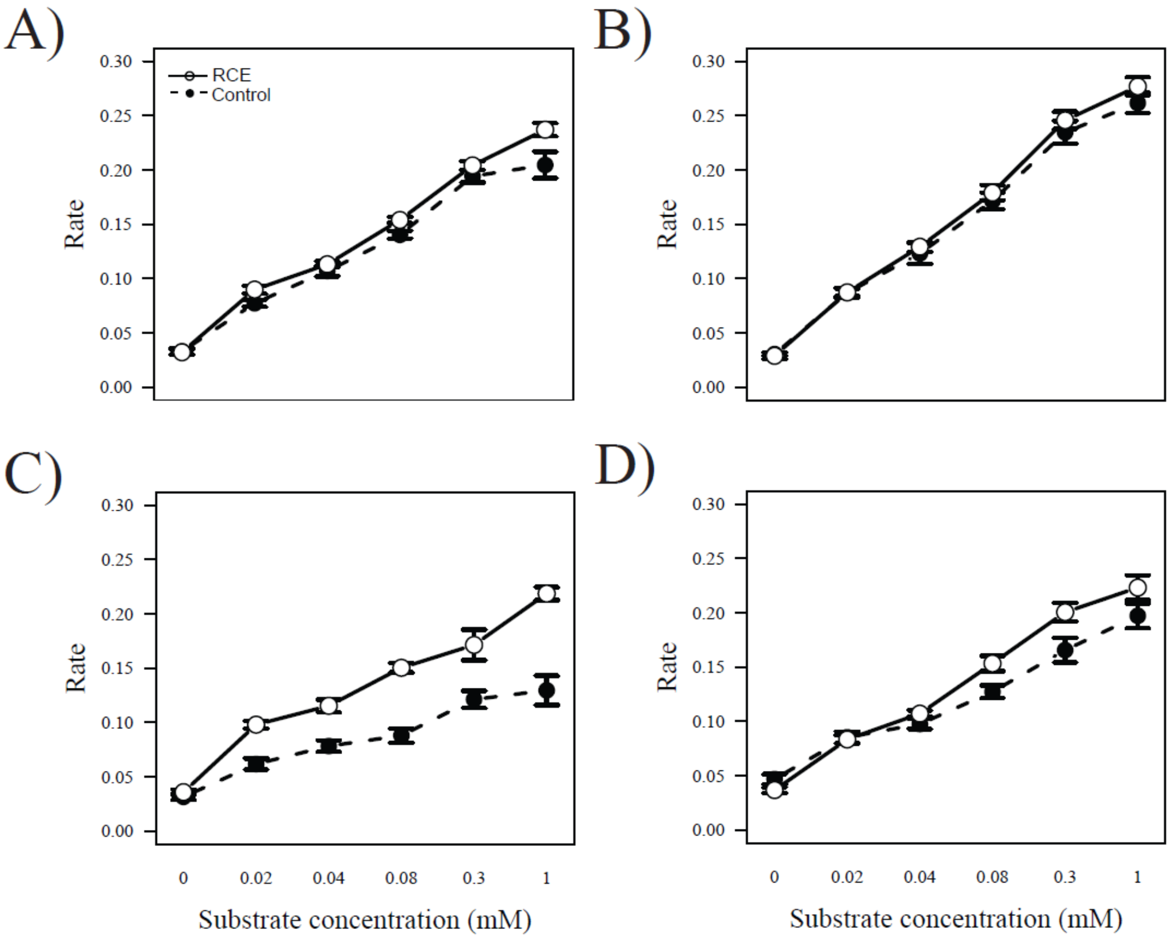


SI Figure 2: Reaction rate (nmol NADPH produced per minute) of glucose-6-phosphate isomerase in Choristoneura fumiferana across populations generally increases following repeated cold exposure (5x -15°C) across substrate concentrations. A) Inuvik population, B) Alberta population, C) New Brunswick population, and D) IPQL strain. N = 5 biological replicates per cold exposure type.

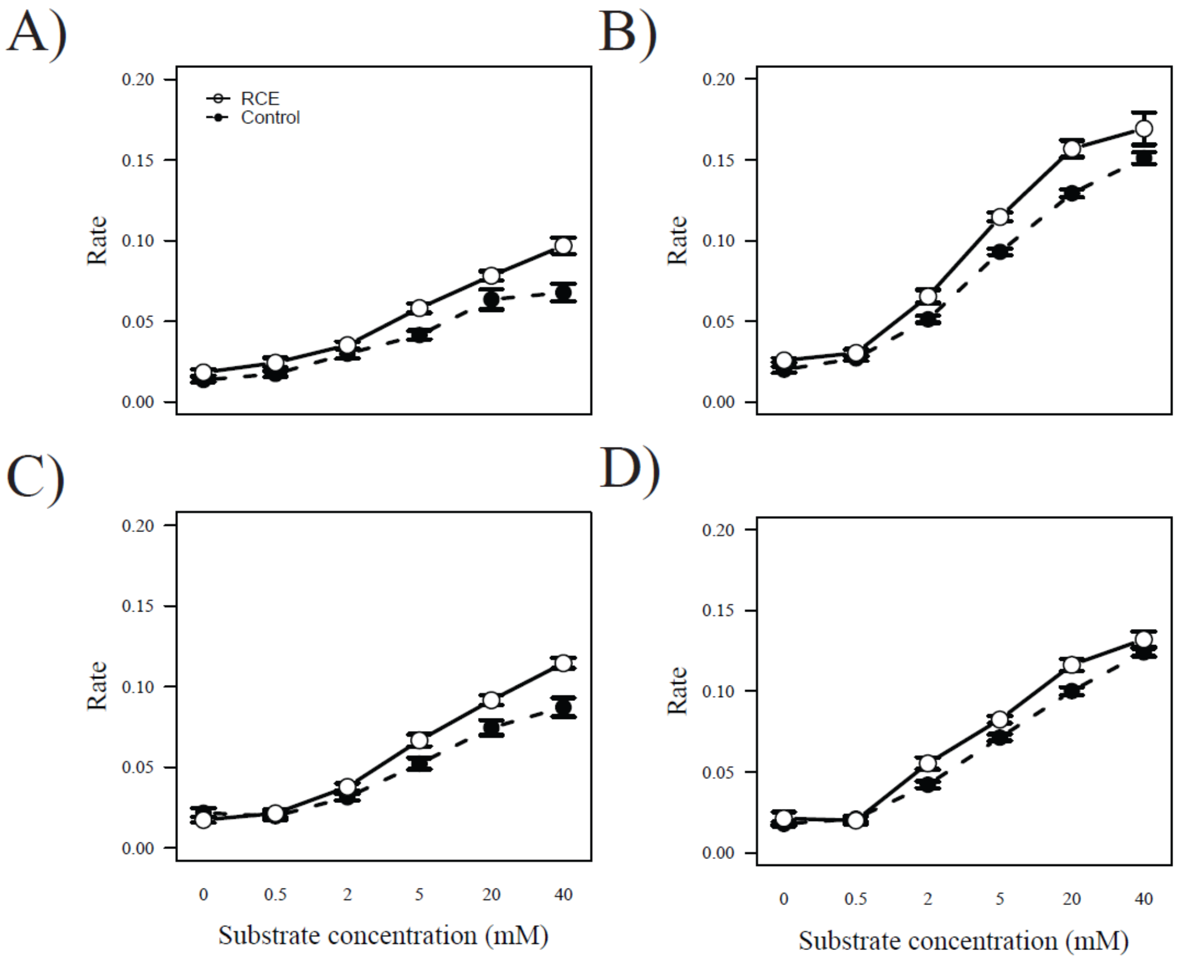


SI Figure 3: Reaction rate (nmol NADPH consumed per minute) of phosphofructokinase in *Choristoneura fumiferana* across populations generally increases following repeated cold exposure (5x -15°C) across substrate concentrations. A) Inuvik population, B) Alberta population, C) New Brunswick population, and D) IPQL strain. N = 5 biological replicates per cold exposure type.

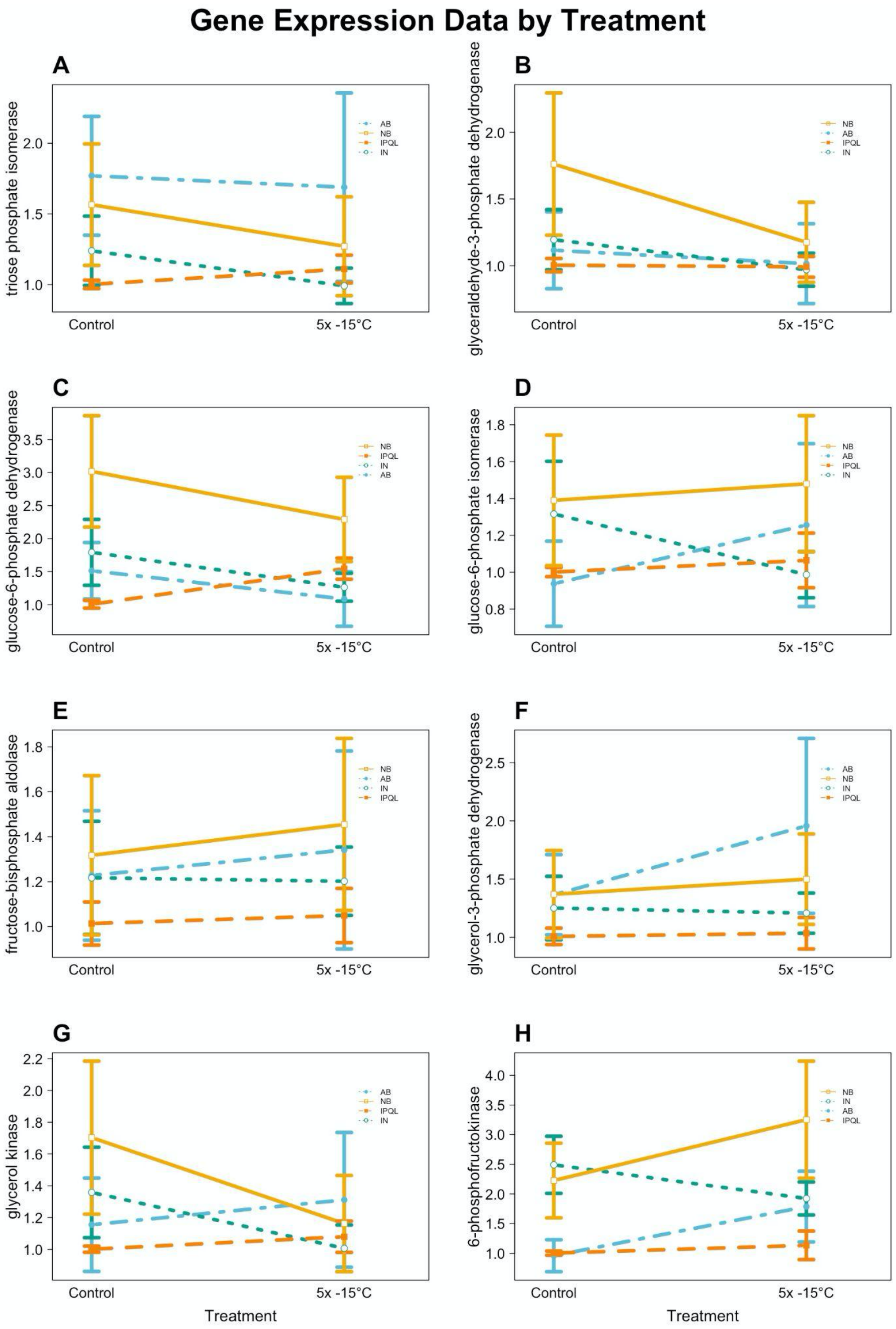


SI Figure 4: Gene expression profiles of eight target genes across different treatment conditions (control and repeated cold exposure of 5x -15°C). Data points represent the mean ± standard error for relative fold change in mRNA abundance. A. Triose Phosphate isomerase. B. Glyceraldehyde 3-phosphate dehydrogenase. C. Glucose-6-phosphate dehydrogenase. D. Glucose-6-phosphate isomerase. E. Fructose-bisphosphate aldolase. F. Glyceraldehyde -3-phosphate. G. Glycerol kinase. H. 6-Phosphate-fructokinase

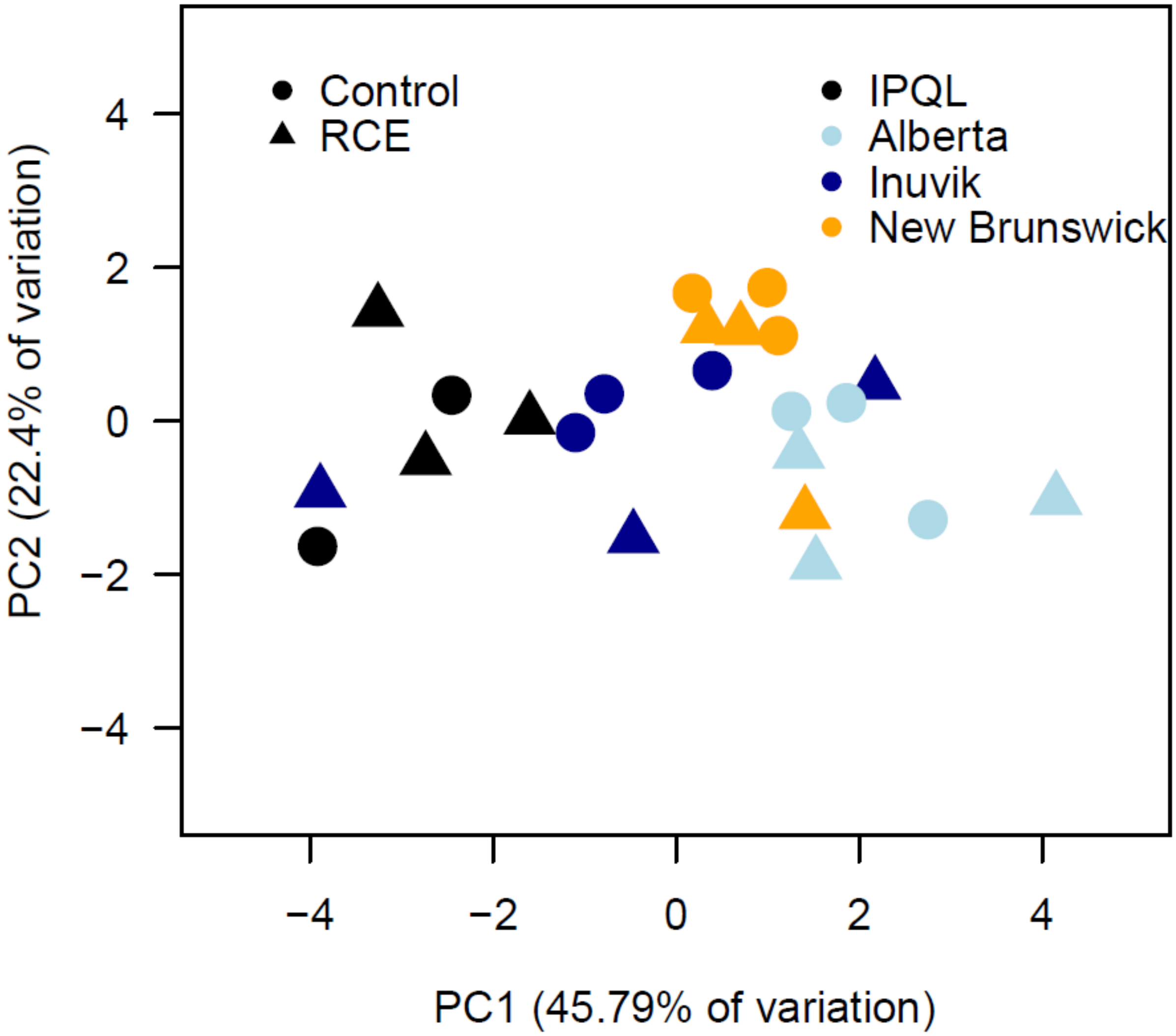


SI Figure 5. Principal components analysis of glycolytic gene expression data from eastern spruce budworm from four populations (“IPQL”, “Alberta”, “New Brunswick”, and “Inuvik”) receiving either control conditions or repeated cold exposures (details in methods).
